# Omnivorous diets of sympatric duck species in a subtropical East Asia wetland unveiled by multi-marker DNA metabarcoding

**DOI:** 10.1101/2025.04.07.647698

**Authors:** Pei-Yu Huang, Emily Shui Kei Poon, Lai Ying Chan, Derek Kong Lam, Ivy Wai Yan So, Yik-Hei Sung, Simon Yung Wa Sin

## Abstract

The East Asian-Australasian Flyway (EAAF) is experiencing notable population decreases in its migratory waterbird species. Our understanding of the foraging ecology of these waterbird species, including ducks, is crucial for monitoring and safeguarding their food sources and wetland habitats. Here, we used a DNA metabarcoding approach to analyze the fecal DNA from duck species to elucidate their dietary composition during the wintering period in a subtropical East Asian wetland. By employing multiple markers (*18S*, *COI*, and *trnL*) targeting different taxonomic groups and levels, we offered a comprehensive dietary analysis for omnivores that consume both plants and animals. We revealed the dietary compositions of common migratory duck species and their intraspecific and interspecific variations. While ducks are generally known to be omnivorous, *Anas crecca* had a more specialized diet and was primarily herbivorous throughout winter. In contrast, the sympatric *Mareca penelope* and *Spatula clypeata* exhibited a more omnivorous foraging behavior. Moreover, *A. crecca* displayed less dietary variation among individuals, while individuals of *M. penelope* and *S. clypeata* were highly variable in their diets. Comparing our results with those of studies conducted in different regions, we found that the dietary compositions of these duck species varied to different degrees across geographic locations. This variation underscores the flexibility of these duck species in their diets and their adaptable foraging strategies. Our findings also indicate that grasslands rich in herbaceous plants and aquatic environments abundant in small aquatic invertebrates are vital foraging habitats for duck species during their winter stay.

## 1. Introduction

Ducks are a diverse group of waterfowl that belong to the family Anatidae, which also includes geese and swans. They are not a monophyletic group and are divided into several subfamilies based on their genetic and physical characteristics (Johnson and Sorenson, 1999). These mostly aquatic birds inhabit both freshwater and saltwater environments, and some duck species, particularly those breeding in the temperate and Arctic regions of the Northern Hemisphere, are known for their long-distance migrations (Arzel et al., 2006). In contrast, ducks living in the tropical areas tend to be non-migratory. Migratory ducks follow through ’flyways’ annually, with the East Asian-Australasian Flyway (EAAF) being one of the most notable, spanning 22 countries and territories. Waterbirds that use the EAAF predominantly breed in far eastern Russia, Alaska, the Mongolian Plateau, and the Amur River basin, and winter in East Asia, Southeast Asia, Australia, and New Zealand (EAAFP, 2024). During migration, waterbirds depend on highly productive wetlands to rest and feed, building up sufficient energy to fuel the next phase of their journey. Ducks, in particular, consume a variety of plants and animals found in wetlands, including grasses, aquatic plants, crustaceans, fish, insects, small amphibians, mollusks, and many other invertebrates (Hitchcock Jr et al., 2021). The abundance, quality, and diversity of these food resources provided by wetlands directly impact the reproductive and survival success of duck species (Holopainen et al., 2015). However, wetlands are increasingly threatened by global changes, and migratory birds in the EAAF are among the world’s most vulnerable to these pressures, given Asia’s large population and booming economies. The EAAF is experiencing significant population declines among its migratory waterbirds (Zhang et al., 2023).

Addressing this issue requires a better understanding of the foraging ecology of ducks and other waterbirds, which is essential for better monitoring and protecting of their wetland habitats for future generations.

Over the last century, considerable effort has been devoted to studying the diets of common duck species in North America and the Western Palearctic (Dessborn et al., 2011, Callicutt et al., 2011). Notably, extensive studies have been carried out on the diets of several duck species such as *Anas acuta*, *Mareca penelope*, *Anas platyrhynchos*, and *Anas crecca* in the Western Palearctic. However, specific knowledge gaps exist in these studies, with geographical and temporal biases identified as key areas of concern (Dessborn et al., 2011). In addition, compared to the North America and Western Palearctic, very few dietary studies on ducks have been conducted in the regions within EAAF. Only China, Japan, South Korea, New Zealand, and Pakistan have conducted such studies, which covered various species, including *Anas* (*A. acuta*, *A. crecca*, *A. chlorotis*, *A. platyrhynchos*, and *A. zonorhyncha*), *Spatula* (*S. querquedula* and *S. clypeata*), *Mareca* (*M. strepera*, *M. penelope*, and *M. falcata*), *Aythya* (*A. fuligula*, *A. ferina*, and *A. nyroca*), and *Hymenolaimus* (*H. malacorhynchos*) (Ando et al., 2023, Shin et al., 2016, Collier, 1991, Raza et al., 2023, Luo et al., 2024).

Studies have shown that food compositions consumed by the same duck species using different flyways varied considerably in relation to the availability of food resources in different parts of their migratory range (Dessborn et al., 2011), it is crucial to conduct more dietary studies on ducks at important breeding and non-breeding locations along EAAF. By doing so, we can better understand the food requirements during the different life cycles of migratory duck species and provide valuable data to protect their critical food sources and habitats.

Hong Kong, a highly developed coastal city in the central part of EAAF, provides essential wetland habitats for migratory waterbirds (Huang et al., 2021). The northwestern region of Hong Kong is a wetland complex that includes natural, semi-natural, and artificial habitats. One vital area of this region is the Mai Po Inner Deep Bay Ramsar Site, which covers approximately 1500 hectares (Huang et al., 2022). Ducks are among the most abundant waterbird groups that winter in the Deep Bay area, with at least thirty species recorded at the site. These include *S. clypeata* (northern shoveler), *A. crecca* (common teal), *M. penelope* (Eurasian wigeon), and *A. acuta* (northern pintail), all of which represent more than 0.25% of the EAAF population (WWFHK, 2024). These four duck species are widely distributed across both the Old and New Worlds, except for *M. penelope*, which is primarily found in the Palearctic range (Kulikova et al., 2019). All these species forage by water dabbling. *Spatula clypeata* is particularly notable for its specialized spatulate bill, uniquely adapted for filtering tiny organisms from the water (Kooloos et al., 1989). However, other species may occasionally engage in similar feeding behavior. *Mareca penelope* sometimes grazes on aquatic vegetation or, like *A. crecca* or *A. acuta*, tipping forward to reach submerged food sources (Ramírez-Albores et al., 2021). Although *S. clypeata*, *M. penelope*, *A. crecca*, and *A. acuta* are classified as ’least concern’ in the IUCN (The International Union for Conservation of Nature) red list (IUCN, 2025), some of their wintering populations have exhibited obvious decline. For example, *M. penelope A. crecca*, *A. acuta* populations declined significantly between 1998 and 2017 (Sung et al., 2021). Their populations are predicted to further decline in the future due to various threats to these duck species, such as wetland habitat loss and avian diseases. Despite the presence of hydrological management in the water ponds in the Ramsar Site (WWFHK, 2023), there is a lack of even fundamental information on food utilization within the managed habitat.

In the past, the primary method used to study the diets of ducks worldwide was through sacrificing them to collect the contents of their esophagus/proventriculus, gizzard or gut for microscopic examination (Miller et al., 2009, Jamieson et al., 2001). These studies have revealed that ducks are omnivores and consume animal and plant matter (Barboza and Jorde, 2001). However, direct examination of digested contents poses many challenges.

Firstly, the taxonomic resolution of digested contents is prone to bias and error, especially for soft-bodied items, which require a high level of taxonomic expertise for proper identification (Nielsen et al., 2018). Secondly, the most common conventional method for quantifying food items in dietary studies is using the frequency of occurrence of and counting the number of items of different taxa. However, relying solely on food item counts may provide little insight into the relative importance of different taxa in terms of nutrition or energy intake (Dessborn et al., 2011). With the recent advent of DNA metabarcoding, it is now possible to collect fecal samples from ducks non-invasively and analyze the composition of fecal DNA in high taxonomic resolution using genetic markers. While it is true that DNA metabarcoding is not a bias-free approach, it has been shown that using relative read abundance information often provides a more accurate view of population-level diet, even with moderate recovery biases incorporated (Deagle et al., 2019). Furthermore, we can use multiple markers in DNA metabarcoding to reveal the relative abundance and frequencies of occurrence of each food item and unveil the relative importance of animal and plant matter in the diet of omnivores (da Silva et al., 2019).

Understanding the foraging ecology of duck species is crucial to their conservation, particularly in regions within EAAF. Therefore, this study addresses important gaps in our understanding of the food use by wintering ducks in Mai Po wetland. Specifically, we use DNA metabarcoding with multiple markers to 1) investigate the dietary compositions of four common migratory duck species, including *S. clypeata*, *A. crecca*, *M. penelope*, and *A. acuta*, to gain insights into the specific food resources required by these duck species wintering in Mai Po wetland, and 2) reveal the intraspecific and interspecific variation in diets among these species. This information will provide critical insights into the foraging strategies of these species and help us better understand how they utilize the wetland habitat during the wintering period.

## 2. Materials and Methods

### 2.1 Sample collection and DNA metabarcoding

The Mai Po Nature Reserve (MPNR; part of the Mai Po Inner Deep Bay Ramsar Site) is a wetland complex comprising five main habitats, including gei wai, freshwater ponds, intertidal mudflats, mangroves, and reedbeds (WWFHK, 2025). Gei wai, or gei wai pond, is a traditional shrimp pond system commonly found in coastal areas of southern China and Hong Kong. These ponds are used for aquaculture, particularly for shrimp farming, and are characterized by a series of interconnected shallow ponds with controlled water flow for raising aquatic species (Cha et al., 1997). We obtained permission to enter the MPNR (22°29’20.7"N 114°02’09.9"E) and collected anatid fecal samples from the ground around gei wai. Samples were collected at dawn, where we specifically gathered fresh feces on the ground near the ponds shortly after the anatids had left (Huang et al., 2021). We spaced the collection points at least 0.5 m apart to avoid duplicating samples from the same individual. Between January and February 2020, 150 fecal samples were collected. These samples were immediately preserved on dry ice in the field and stored at -80 °C until DNA extraction (Supplementary Materials and Methods) (Huang et al., 2022). DNA barcoding was conducted on each sample collected to identify host species (Supplementary Materials and Methods). Specifically, we obtained 57 samples from *M. penelope*, 48 samples from *S. clypeata*, 42 from *A. crecca*, and three from *A. acuta*. Our analysis included mock communities (Table S1, Supplementary Materials and Methods) and negative controls during the DNA extraction process to ensure accuracy. We used the Qubit dsDNA high-sensitivity (HS) assay on a Qubit 4 fluorometer (Invitrogen, Carlsbad, USA) for DNA quantification. To identify the dietary composition, we employed specific genetic markers tailored to the dietary habits of anatids, known to be omnivorous. The first marker used was a universal genetic marker (forward: 5’GGTCTGTGATGCCCTTAGATG3’ and reverse: 5’GGTGTGTACAAAGGGCAGGG3’) that amplifies the V7 region of the 18S small subunit of nuclear ribosomal DNA (rDNA; ca. 170 bp) (McInnes et al., 2017). Additionally, we utilized a COI marker (forward: 5’CCIGAYATRGCITTYCCICG3’ and reverse: 5’GGIGGRTAIACIGTTCAICC3’) that specifically targets the Folmer region of the mitochondrial cytochrome c oxidase I (COI; ca. 86 bp). This marker is particularly effective in identifying invertebrates (Elbrecht and Leese, 2017). Lastly, a plant-specific marker (forward: 5’GGGCAATCCTGAGCCAA3’ and reverse: 5’CCATTGAGTCTCTGCACCTATC3’) was employed to amplify a variable region of the P6 loop in the chloroplast *trnL* (UAA; ca. 10-143 bp) intron (Taberlet et al., 2007). Fecal DNA samples, mock communities, and negative controls were used for library preparation through a two-step polymerase chain reaction (PCR) process (Wan et al., 2024, Huang et al., 2021, Huang et al., 2022, Wei et al., 2024) with the *18S*, *COI*, and *tnrL* markers (Supplementary Materials and Methods). To create a library multiplex, we combined the individual libraries in an equimolar ratio for each marker. Each multiplex was sequenced to a depth of approximately 400 k reads on a NovaSeq instrument (PE 150 bp reads) by the Novogene Corporation (Hong Kong).

### 2.2 Bioinformatics

The demultiplexed raw paired-end fastq reads of *18S*, *trnL*, and *COI* markers were preprocessed by paired read merging, adapter trimming, and quality filtering. The paired-end reads were merged using USEARCH v11.0.667 with the -fastq_mergepairs function (Edgar, 2010). PCR primer sequences were trimmed using CUTADAPT v2.4 with the linked adapter mode (max_error_rate=0.15) (Martin, 2011). Only the merged reads that matched the primer sequences for the *18S*, *trnL*, or *COI* markers were retained. The quality of reads was assessed using FastQC v0.11.8 (Wingett and Andrews, 2018) and the -fastq_eestats2 command in VSEARCH (Rognes et al., 2016). Subsequently, we retained the high-quality reads that fell within the target lengths (*18S*: 130−230 bp; *trnL*: 10-100; *COI*: 75−85 bp). These reads also had an expected number of errors per read < 1 (-fastq_maxee 1), which was determined using the -fastq_filter function in VSEARCH. All preprocessed reads were dereplicated using the VSEARCH -derep_fulllength. From the dereplicated reads, we generated amplicon sequence variants (ASVs) by removing the chimeras and singletons (with abundance < 0.0001% of all reads) using the USEARCH -unoise3 (Edgar, 2016b). To cluster all the preprocessed reads into ASVs, we used a similarity threshold of 99% for *18S* and *trnL*, and 95% for *COI* (VSEARCH -usearch_global -id 0.99/0.95).

The taxonomic classification process was conducted in two steps to achieve a higher taxonomic resolution. Firstly, each ASV was assigned to the lowest identifiable taxonomic level using the SINTAX algorithm in USEARCH (Edgar, 2016a) with a bootstrap cutoff of 0.7. For the *18S* dataset, the ribosomal RNA database SILVA (Glöckner et al., 2017) was utilized. The *trnL* dataset was classified using the CRUX database from the Anacapa Toolkit (Curd et al., 2019), while the *COI* dataset was classified using the mitochondrial database MIDORI (Leray et al., 2022). Secondly, we searched the ASVs from these datasets against the NCBI nt database (non-redundant nucleotide sequences) (Sayers et al., 2022). We extracted the top 1,000 blast hits that exhibited similarity above 90% and an e-value < 1e-50 for *18S*, above 90% and an e-value < 1e-5 for *trnL*, and above 80% and an e-value < 1e-5 for *COI*. Afterward, we assigned the lowest common taxonomic level shared by 95% of *18S* blast hits (≥ 100 bps), 80% of *trnL* blast hits (≥ 40 bps), and 80% of *COI* blast hits (≥ 50 bps) using BASTA (Kahlke and Ralph, 2019) with the lowest common ancestor (LCA) algorithm. The results were then combined to assign ASVs with lower ranks of taxonomies.

To eliminate potential false-positive reads and contaminant ASVs, we applied thresholds defined by mock communities. We also removed non-dietary items (such as Humans, Aves, Bacteria, and Protists) and those with low taxonomic resolution (such as Eukaryota and Chordata). After these treatments, we removed samples with low read numbers, guided by the read number-based rarefaction curves. We obtained a mean of 237,059 reads per sample for *18S*, 227,500 reads per sample for *trnL*, and 97,084 reads per sample for *COI*.

### 2.3 Data analysis

We used 132 samples from four anatid species for downstream analysis, including *S. clypeata* (n=44 for *18S*, n=42 for *trnL*, and n=45 for *COI*), *M. penelope* (n=41 for *18S* and *COI*; n=42 for *trnL*), *A. crecca* (n=41 for *18S* and *COI*; n=42 for *trnL*), and *A. acuta* (n=3 for *18S* and *trnL*, n=2 for *COI*). For the *18S* dataset, we identified 39 taxonomic categories from 73 taxa represented by 129 ASVs. The *trnL* dataset yielded 47 taxonomic categories from 73 taxa represented by 125 ASVs. The *COI* dataset resulted in 27 taxonomic categories from 132 taxa represented by 514 ASVs. To present the data, we calculated three metrics: (i) relative read abundance (RRA), which represents the percentage of read count for each taxon in a sample; (ii) weighted percentage of occurrence (wPOO), which indicates the percentage of occurrence for each taxon in a sample, and (iii) frequency of occurrence (FOO), which measures the proportion of samples in which a taxon is detected (Lee et al., 2021). The RRA or wPOO at the population level is presented as the mean of RRA or wPOO of all individual samples of an anatid species. FOO estimates the frequency of incidence of a taxon within all samples of an anatid species. The results were visualized with the R packages ggplot2 v3.3.5 (Hadley, 2016).

#### 2.3.1 Diversity analysis on the diet of three anatid species

As the number of samples for *A. acuta* was small, they were excluded from the diversity analysis. We utilized Hill numbers (represented as ’D’) (Hill, 1973) for the diversity analyses using the R package hilldiv v1.5.1 (Alberdi and Gilbert, 2019). The sensitivity of Hill number towards abundant ASVs can be modulated by the *q* value (the order of diversity). When the q value increases, more weight is given to the abundant ASVs. The richness of the dietary compositions is indicated by the Hill number of order *q*=0 (^0^D), which considers only the occurrence of each ASV. The Hill number of order *q*=1 (^1^D) considers both richness and evenness, which is equivalent to the exponential of Shannon’s diversity index. For the Hill number of order *q*=2 (^2^D), abundant ASVs are given more weight, resulting in a value that corresponds to the multiplicative inverse of Simpson’s dominance index (Jost, 2006).

Hill numbers can be used to calculate alpha, beta, and gamma diversities, which are defined as ^q^D_γ_ = ^q^D_α_ × ^q^D_β_. Alpha diversity (^q^D_α_) measures diversity at the sample level, while gamma diversity (^q^D_γ_) represents diversity at the population or species level. These diversities can be computed using the function div_profile in hilldiv. Beta diversity (^q^D_β_) measures the differences between individual samples and is derived by dividing gamma diversity by alpha diversity (Chao et al., 2012, Jost, 2007). We conducted pairwise comparisons using the Kruskal-Wallis test to compare the dietary diversity between anatid species at the sample level (alpha diversity). We then performed a post hoc Dunn test, using Benjamini-Hochberg correction (*p*<0.05) with the div_test function of the hilldiv package.

To assess the differences in taxa compositions between anatid species, we calculated pairwise binary Jaccard dissimilarity distances based on the occurrence of each ASV, as well as pairwise Bray‒Curtis dissimilarity distances based on the fourth root transformed RRA of each ASV. The results were visualized using Principal Coordinates Analysis (PCoA) with the ordinate and plot_ordination functions in the R package phyloseq v1.30.0 (McMurdie and Holmes, 2019). We performed hierarchical clustering analysis using Ward’s method in the hclust function. Additionally, we conducted a Permutational Multivariate Analysis of Variance (PERMANOVA) test to evaluate the separation of dietary compositions between the anatid species. This test evaluated the centroid and dispersion of diet compositions for individual samples within each group in a measure of space. To ensure the significance of interspecific variation in the PERMANOVA test, we checked for homogeneity of intra-group beta-dispersion (p>0.05) using the adonis function in the vegan v2.5.7 package (Oksanen et al., 2019). We conducted pairwise PERMANOVAs of the anatid species using the pairwise adonis function (Martinez Arbizu, 2020) and evaluated their beta-dispersions using the betadisper function in the vegan package. To determine the contribution of individual taxa to the variations between anatid species, we calculated the Similarity Percentage (SIMPER). The SIMPER analysis used the non-parametric Kruskal‒Wallis rank-sum test by the simper.pretty and kruskal.pretty functions in R scripts simper_pretty.R and R_krusk.R (Steinberger, 2020). Only taxa that exhibited statistically significant variance (*p*<0.05) were presented. PERMANOVAs and SIMPER analyses based on the occurrence data were conducted using binary Jaccard dissimilarity distances, while those based on RRA were performed using Bray‒Curtis distances.

## 3. Results

### 3.1 Dietary compositions of A. crecca, M. penelope, S. clypeata, and A. acuta

Based on the *18S* data, it was found that *A. crecca* exhibited the highest and the most frequent consumption of streptophytes (94%RRA and 66%wPOO) among the four species studied (Fig. 1; Tables S2-4). The results also revealed that *A. crecca* consumed tiny proportions of algae, fungi, and arthropods, including species in Tenuipalpidae (false spider mites), Neoptera, Diptera (flies), as well as crustaceans, such as species in Podocopida and copepods *Nannopus*. Further analysis of the *trnL* data revealed that the streptophytes targeted by *A. crecca* were primarily asters from the Asteraceae family (83%RRA and 40%wPOO among plants), as well as monocotyledonous grasses from the Poaceae family in the order Poales (8%RRA and 42%wPOO among plants), such as common reed (*Phragmites australis*). A tiny proportion of Myrtales and various flowering plants, including those from the Fabales, Fagales, Malpighiales, and Caryophyllales orders, were also consumed by *A. crecca* (Fig. 1; Tables S5-7). Additionally, the analysis of *COI* data showed that *A. crecca* consumed arthropods (90%RRA and 78%wPOO among marcroinvertebrates), including dipterans (e.g., typical mosquitoes in *Culex*), lepidopterans (butterflies and moths), coleopterans (beetles), and arachnids. It also fed on small proportions of sponges, gastropods, and annelids (Fig. 1; Tables S8-10).

**Figure 1.**
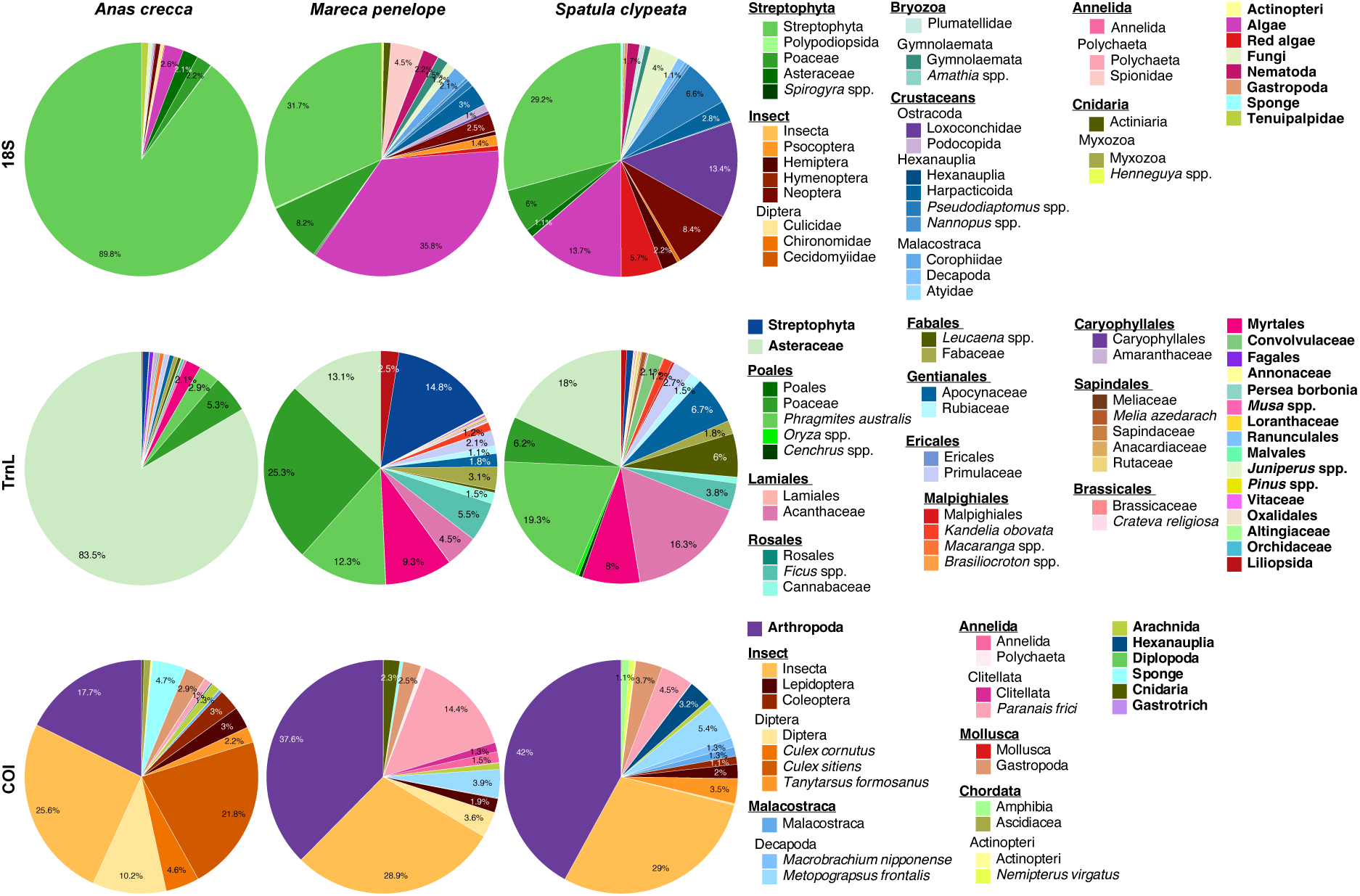
Dietary compositions of three wintering anatid species were determined using *18S* rDNA, *trnL*, and *COI* markers on fecal DNA. We used samples from *Spatula clypeata* (n=44 for *18S*, n=42 for *trnL*, and n=45 for *COI*), *Mareca penelope* (n=41 for *18S* and *COI*; n=42 for *trnL*), and *Anas crecca* (n=41 for *18S* and *COI*; n=42 for *trnL*). Each taxon was categorized to the lowest taxonomic level when the number of reads was >0.1% of all taxa detected. Otherwise, the taxa were grouped into higher levels of taxonomic category. The relative read abundance (RRA) of dietary categories are shown as color blocks. Only categories with an RRA >1% are indicated (see Tables S2, S5, and S8 for details).

Regarding *M. penelope*, the analysis of *18S* data indicated that it had a heavy and frequent consumption of streptophytes (40%RRA and 44%wPOO), particularly Poaceae (Fig. 1; Tables S2-4). Algae also comprised a considerable portion of its diet (36%RRA and 25%wPOO). While arthropods (12%RRA and 13%wPOO), polychaetes (bristle worms, e.g., spionids) in the Annelida phylum, as well as species in Nematoda (roundworms) and Bryozoa (moss animals, mainly Gymnolaemata) were preyed upon, they constituted smaller proportions of *M. penelope*’s diet. The results from *trnL* data exhibited a similar pattern to the *18S* data for plant consumption. It showed that a high consumption of grass species from the Poaceae family (38%RRA and 24%wPOO among plants), such as common reed (*P. australis*) (Fig. 1; Tables S5-7). *M. penelope* also ate plant parts from various orders such as Myrtales, Rosales (e.g., *Ficus* figs and Cannabaceae), Lamiales (e.g., Acanthaceae), Fabales, Gentianales (Apocynaceae and Rubiaceae), Ericales (Primulaceae), Malpighiales (e.g, *Kandelia obovata*) orders, among others. Results from *COI* data indicated that arthropods (77%RRA and 70%wPOO among invertebrates) make up the majority of *M. penelope*’s invertebrate diet, with insects (34%RRA and 34%wPOO among invertebrates), such as dipterans and lepidopterans, being the most frequently consumed (Fig. 1; Tables S8-10). Decapods, including marsh crabs like *Metopograpsus frontalis*, were also preyed upon. *Mareca penelope* also fed on smaller proportions of annelids (18%RRA and 20%wPOO among invertebrates), mainly clitellate detritus worms like *Paranais frici*, as well as gastropods (slugs and snails) and cnidarians.

The *18S* data revealed that the diet of *S. clypeata* primarily consisted of streptophytes and arthropods (RRA: 36% and 36%; wPOO: 39% and 25%), and the later was higher than the other anatids. There were notable contributions from two plant taxa, such as Poaceae and Asteraceae (Fig. 1; Tables S2-4). The consumption of arthropods in *S. clypeata* included crustaceans, such as ostracods (13%RRA and 5%wPOO, e.g., loxoconchidae), Hexanauplia (10%RRA and 12%wPOO, e.g. the order Harpacticoida, *Pseudodiaptomus* spp., and *Nannopus* spp.), and insects (11%RRA and 7%wPOO. e.g.

Neoptera and Hemiptera). Other dietary components included algae (14%RRA and 15%wPOO), red algae (6%RRA and 9% wPOO). Consistent with *18S* data, the *trnL* data showed that *S. clypeata* primarily consumed plant taxa in Poaceae (27%RRA and 18%wPOO, e.g., *P. australis*) and Asteraceae (18%RRA and 9%wPOO). Additionally, abundant Lamiales were detected in *trnL* data (16%RRA and 8%wPOO, e.g., Acanthaceae) (Fig. 1; Tables S5-7). *Spatula clypeata* also fed on plant materials from a diverse range of taxa, including species in Gentianales (Apocynaceae and Rubiaceae), Myrtales, Fabaceae, Rosales (e.g., figs *Ficus* in Moraceae), Ericales (e.g., Primulaceae), Malpighiales (such as *Kandelia obovata*), Convolvulaceae (bindweeds), Sapindales, and more. Regarding invertebrates, arthropods (90%RRA and 81%wPOO) formed the main prey for *S. clypeata* (Fig. 1; Tables S8-10), including insect species (36%RRA and 35%wPOO) such as dipterans (e.g., lake flies *Tanytarsus formosanus*), lepidopterans, and coleopterans. They also consumed crustaceans (11%RRA and 11%wPOO) such as malacostracans (e.g., oriental river prawn *Macrobrachium nipponense* and marsh crabs *Metopograpsus frontalis*) and species in Hexanauplia. The diet also included small proportions of annelids (e.g., *P. frici*), gastropods, and amphibians.

Despite its small sample size, *A. acuta* predominantly fed on streptophytes (62%RRA and 48%wPOO in *18S* data), mainly grasses from Poaceae (Tables S2-4). Additionally, it fed on arthropods (32%RRA and 22%wPOO in *18S* data), especially benthic copepods of the Harpacticoida order, planktonic copepods in *Pseudodiaptomus*, ostracods in the subclass Podocopa, and algae (Fig. 1; Tables S2-4 and S8-10). *TrnL* data revealed more plant species being consumed, in which asters from Asteraceae (30%RRA and 11%wPOO), bindweeds in Convolvulaceae (27%RRA and 11%wPOO) and grasses in Poaceae (13%RRA and 23%wPOO) are some of the primary plants (Fig. 1; Tables S5-7). It also feds on other plants in small amounts, including Acanthaceae, Apocynaceae, and Rubiaceae. Arthropoda, especially insects, was the only invertebrate detected in the diet of *A. acuta* as revealed from the *COI* data.

Our findings identified several food taxa commonly found in the diets of all or most of the anatid species studied. These include Asteraceae, species from Poaceae (such as *P. australis*), as well as members of Acanthaceae, Myrtales, Apocynaceae, algae, Podocopida (e.g., Loxoconchidae), red algae, fungi, Nematoda, and various insects. (Fig. S1, Tables S2-S10).

### 3.2 Intraspecific and interspecific variation in anatid species

The analysis of the three genetic markers did not reveal obvious intraspecific variation within *A. crecca* (*A. acuta* was excluded here because of its small sample size). *M. penelope* and *S. clypeata* displayed higher levels of intraspecific variation than those observed in *A. crecca* (Fig. 2 and S2; Tables S11-16).

**Figure 2.**
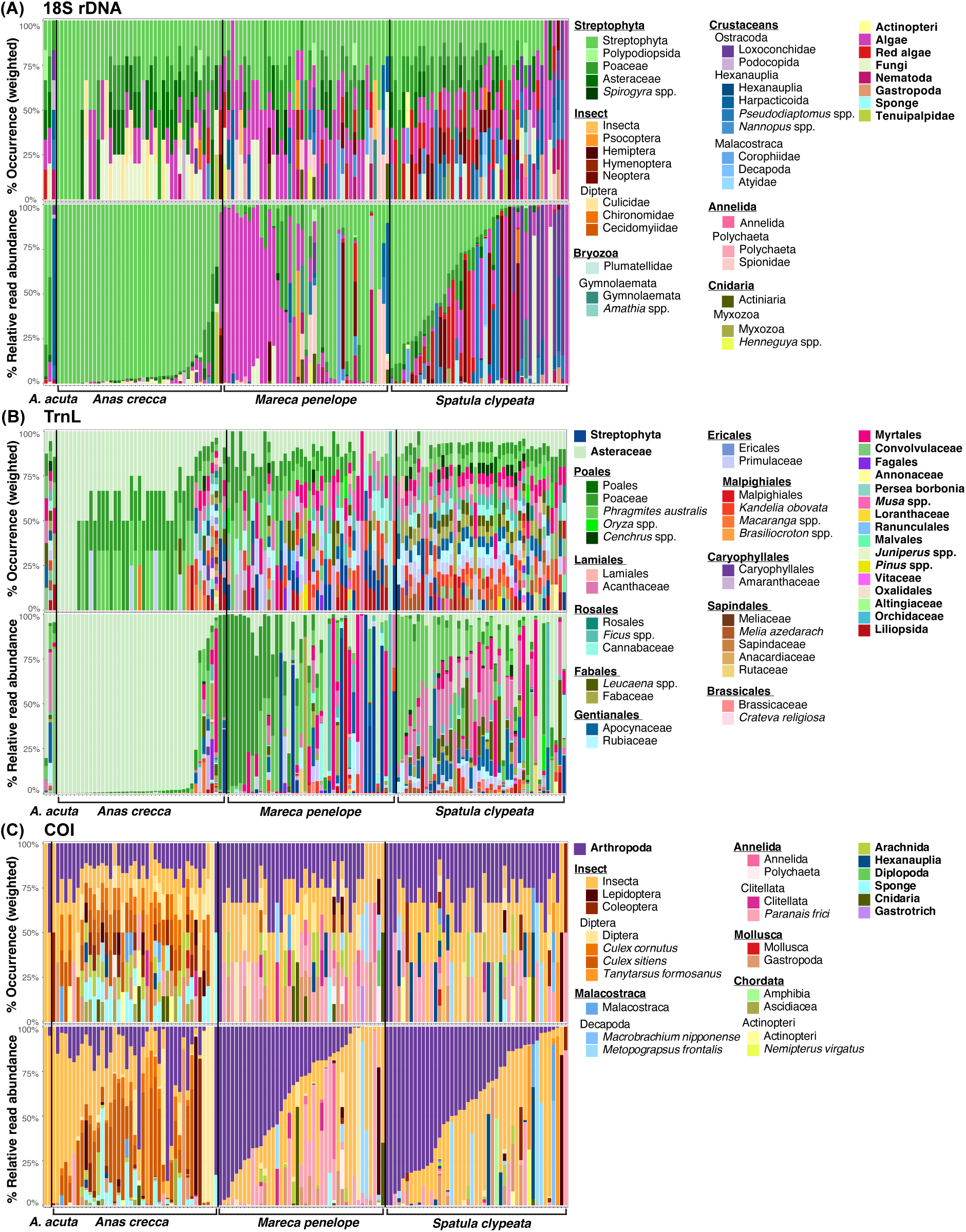
Dietary compositions of individual fecal samples of wintering anatids were detected using (A) *18S* rDNA, (B) *trnL*, and (C) *COI* markers. We used samples from *Spatula clypeata* (n=44 for *18S*, n=42 for *trnL*, and n=45 for *COI*), *Mareca penelope* (n=41 for *18S* and *COI*; n=42 for *trnL*), *Anas crecca* (n=41 for *18S* and *COI*; n=42 for *trnL*), and *Anas acuta* (n=3 for *18S* and *trnL*, n=2 for *COI*). The weighted percentage of occurrence and relative read abundance of each taxon are shown, with each colored bar representing one anatid individual (see Tables S11-16 for details).

Analysis of the alpha diversities showed that, at the individual level, *S. clypeata* had the highest diversities in their plant (*trnL*) diet, followed by *M. penelope* and *A. crecca* (Fig. 3 and 4; Tables S17-18). Despite having the highest plant diversity at the individual level, the diversities of invertebrate (*COI*) in the diets of *S. clypeata* individuals were the lowest among the anatid species based on taxa richness (Fig. 3 and 4; Tables S17-19). *Anas crecca* displayed the highest invertebrate diversities in individual diets. However, we observed decreasing trends in plant and invertebrate diversity values as *q* values increased in all three anatid species. This indicated that individual diets were dominated by several ASVs, and the consumption of plant and invertebrate taxa was uneven in these anatid individuals (Fig. 4).

**Figure 3.**
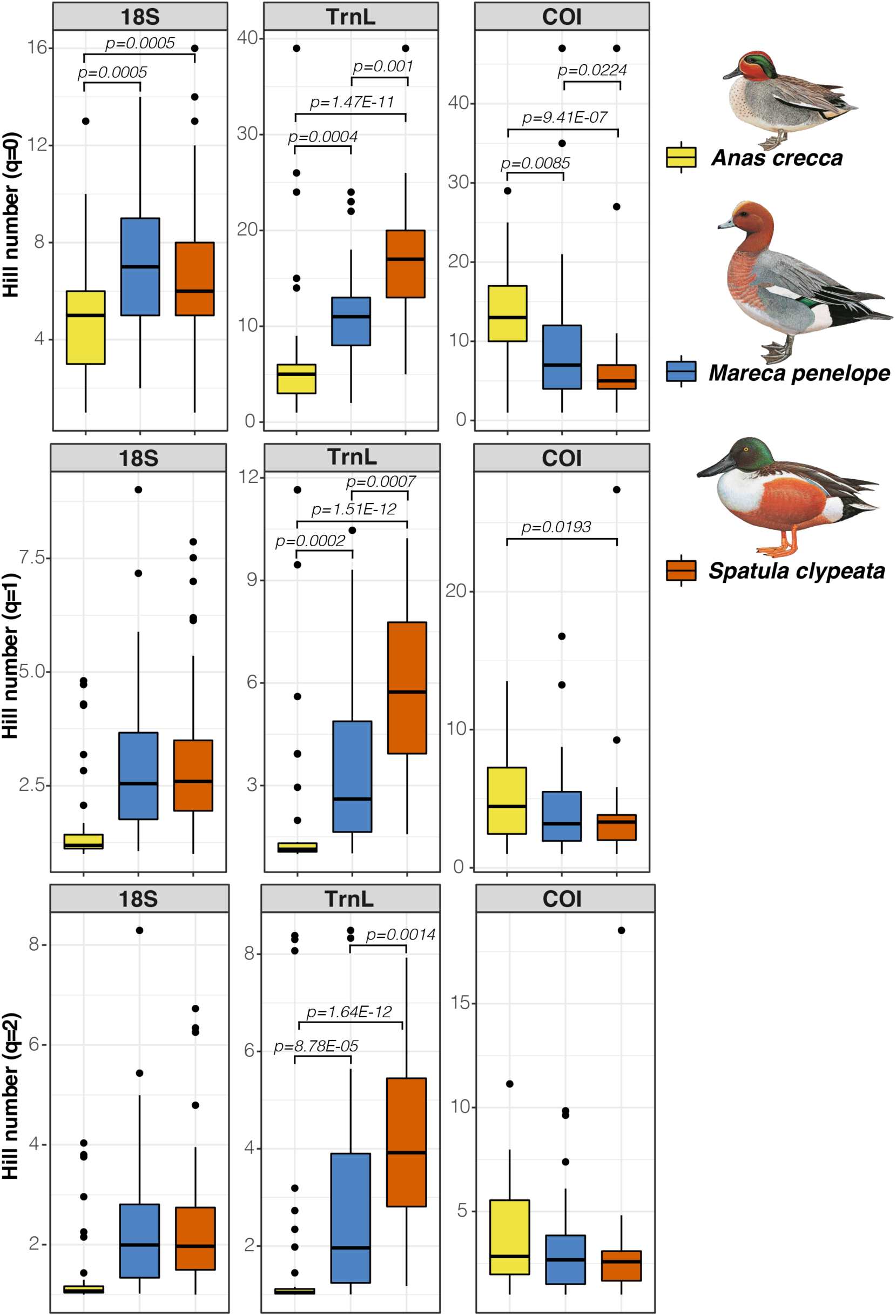
Alpha diversities of the dietary compositions in three species of anatid individuals, as characterized by *18S* rDNA, trnL, and COI markers. We analyzed fecal samples from *Spatula clypeata* (n=44 for *18S*, n=42 for *trnL*, and n=45 for *COI*), *Mareca penelope* (n=41 for *18S* and *COI*; n=42 for *trnL*), and *Anas crecca* (n=41 for *18S* and *COI*; n=42 for *trnL*). The Hill numbers were calculated for three levels of diversity (*q*=0, 1, and 2), with increasing weight given to the abundance of dietary taxa. Each colored box represents the interquartile range, with the median indicated by a line. The whiskers extend to the highest and the lowest values within the 1.5× interquartile range, and the black dots represent outliers. The *p*-values of significant differences between groups are shown above the boxes (see Table S18 for details). Illustrations of anatids were reproduced with the permission of Lynx Edicions.

**Figure 4.**
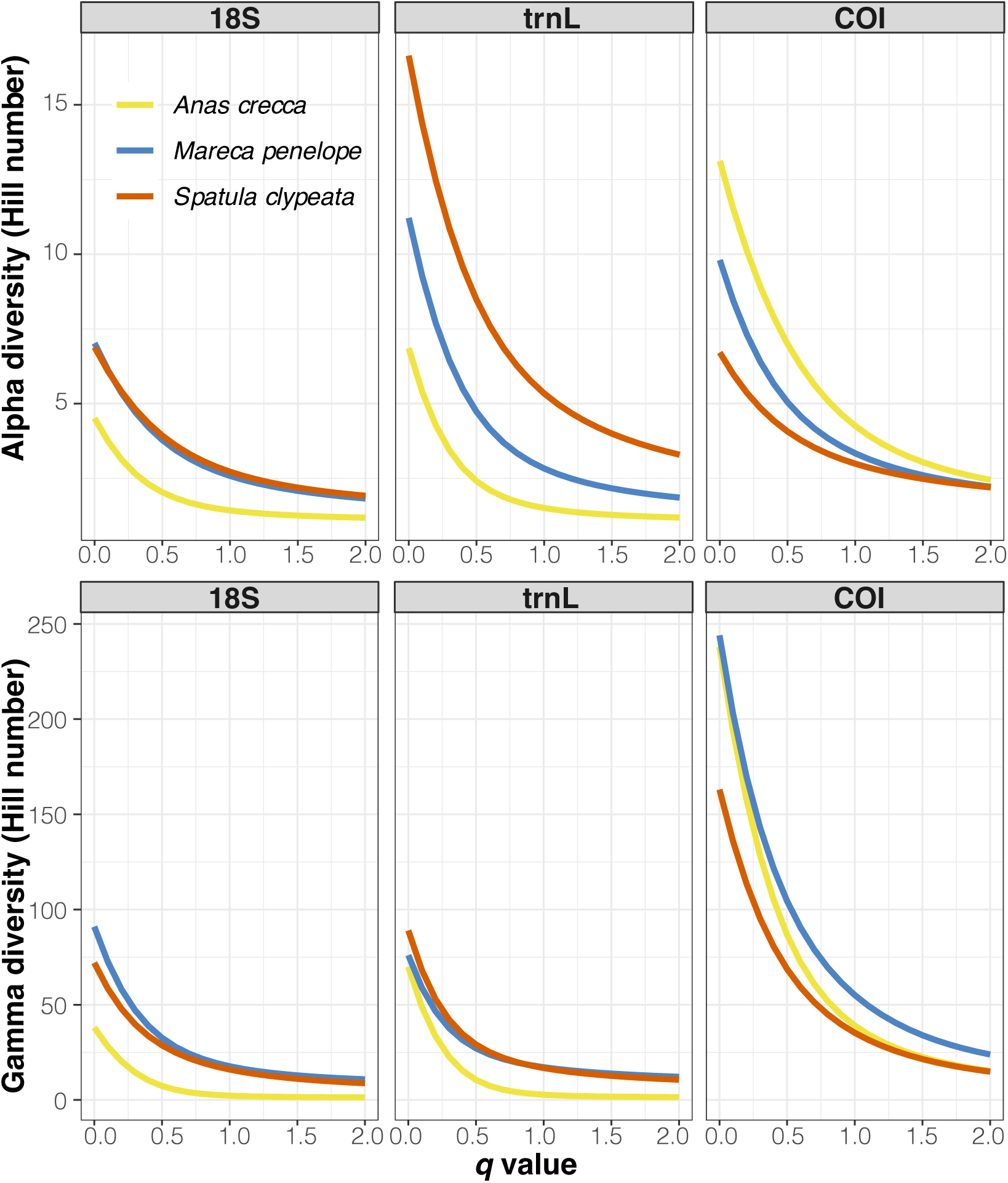
Diversity profiles of the dietary compositions in three anatid species, characterized by *18S* rDNA, *trnL*, and *COI* markers. We analyzed fecal samples from *Spatula clypeata* (n=44 for *18S*, n=42 for *trnL*, and n=45 for *COI*), *Mareca penelope* (n=41 for *18S* and *COI*; n=42 for *trnL*), and *Anas crecca* (n=41 for *18S* and *COI*; n=42 for *trnL*). The alpha and gamma diversities are presented as Hill numbers, with increasing orders of diversity *q* (see Table S17 for details).

At the population level, analysis on gamma diversities based on *18S* (overall) and *trnL* (plant) data showed that *S. clypeata* and *M. penelope* had similar levels of diversities in their diets, whereas the dietary diversities of *A. crecca* were the most unevenly distributed and the lowest among the three species (Fig. 4 and Table S17). Although the diets of *M. penelope* and *A. crecca* shared similar invertebrate (*COI*) taxa richness at the population level, the dietary taxa in *A. crecca* were more unevenly distributed than *M. penelope* and the invertebrate diversity of *A. crecca* became more similar to that of *S. clypeata* as *q* values increased.

The dietary niche of *M. penelope* and *S. clypeata* largely overlapped, whereas *A. crecca* had a more distinct dietary composition, revealed by the three genetic markers based on both Jaccard and Bray–Curtis dissimilarities (Fig. 1, 2, 5, S2). All plant and invertebrate diets were significantly segregated among three anatid species (PERMANOVA, *p-value* =0.001) (Table S19). However, homogeneous intraspecific dispersion was only detected in the abundance-based invertebrate diets (*COI* beta dispersion = 0.07), indicating that the significant separation of overall and 18S diets might be due to the intraspecific heterogeneity rather than interspecific variations.

**Figure 5.**
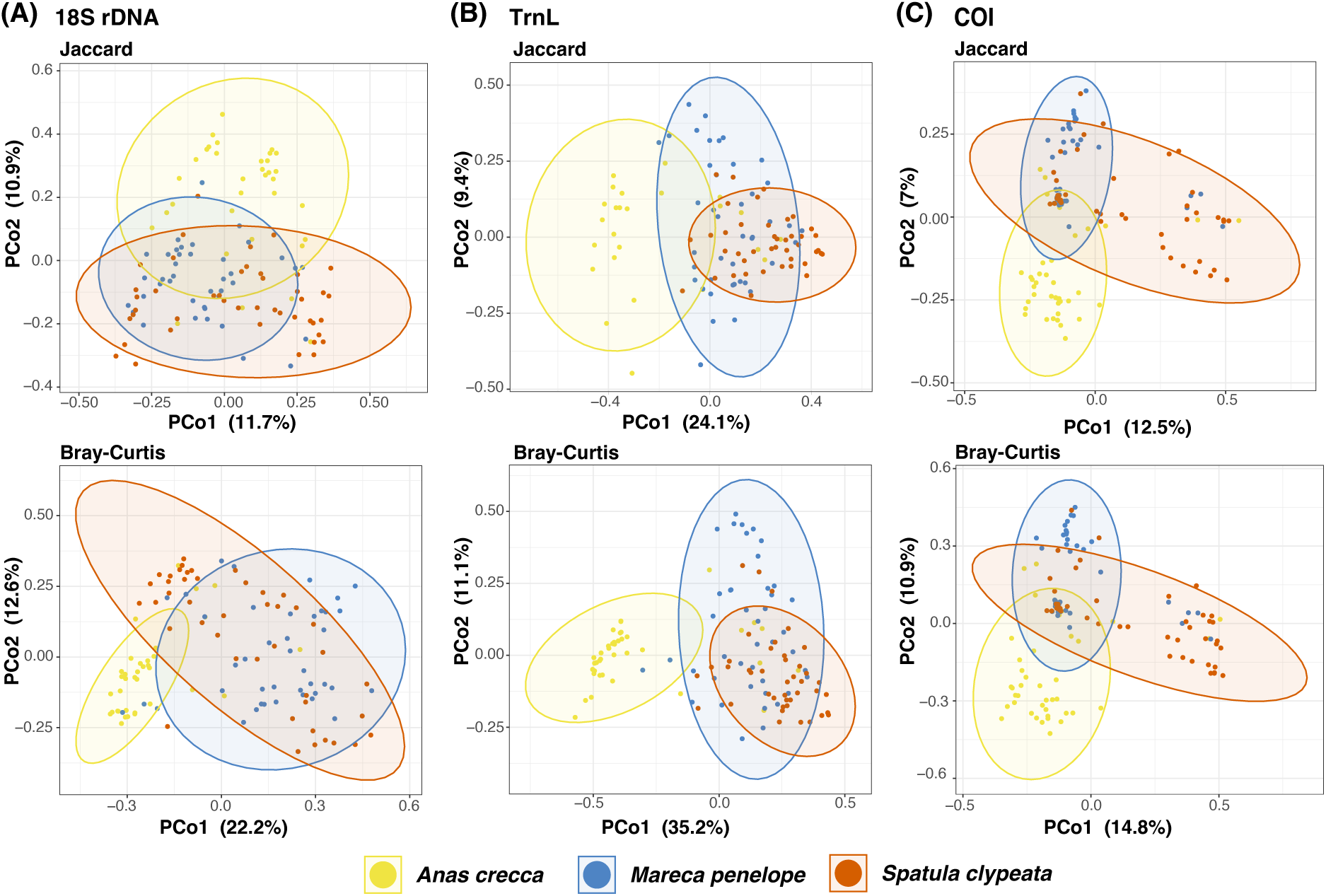
Principal coordinate analysis (PCoA) of dietary compositions in three anatid species, detected using (A) *18S* rDNA, (B) *trnL*, and (C) *COI* markers. We analyzed fecal samples from *Spatula clypeata* (n=44 for *18S*, n=42 for *trnL*, and n=45 for *COI*), *Mareca penelope* (n=41 for *18S* and *COI*; n=42 for *trnL*), and *Anas crecca* (n=41 for *18S* and *COI*; n=42 for *trnL*). The analysis is based on Bray–Curtis distances estimated from relative read abundance data and binary Jaccard distances calculated from occurrence data. The percentages of variations in diet compositions are shown in brackets along axes (see Tables S19-22 for details).

According to the SIMPER results, a group of food taxa contributed to the dietary difference between *A. crecca* and the other two anatid species (Tables S20-22). For example, *A. crecca* consumed higher proportion of streptophytes, including asters from Asteraceae (Tables S5-7), dipterans like typical mosquitoes, such as *Culex* spp., and sponge (Tables S8-10), while consuming fewer items in Poaceae, particularly *P. australis*, Myrtales, Acanthaceae (Tables S5-7), algae (Tables S2-4), clitellate oligochaete worm, *P. frici* (Tables S8-10), compared to *M. penelope* and *S. clypeata*. Although the diets of *M. penelope* and *S. clypeata* were similar to each other, they were slightly differentiated by the consumption of a few taxa (Table S21). For instance, plants in Asteraceae, Acanthaceae, *P. australis*, Apocynaceae (Tables S5-7), as well as red algae, copepods in *Pseudodiaptomus*, insects in Neoptera, and ostracods in Loxoconchidae (Tables S2-4) were consumed more by *S. clypeata* than *M. penelope*. In contrast, plants in Poaceae (Tables S5-7) and algae (Tables S2-4 and S8-10) were consumed more by *M. penelope* compared to *S. clypeata*.

## 4. Discussion

While ducks are generally considered omnivorous, certain duck species, such as *A. crecca* in Hong Kong, demonstrate distinct dietary niches when compared to their sympatric counterparts. The analysis using the three markers indicates that *M. penelope* and *S. clypeata* exhibited omnivorous foraging behavior during their winter stay in Hong Kong. In contrast, although *A. crecca*’s diet included some prey taxa such as insects and gastropods, it was predominantly herbivorous throughout its wintering period in Hong Kong. Additionally, *A. crecca* exhibited less variation in diets between individuals, while individuals of *S. clypeata* exhibited more variations in their overall and animal diets. *Anas crecca* showed lower diversity in its plant diet but higher diversity in its animal diet. Conversely, *S. clypeata* exhibited the highest diversity in its plant diet and the lowest diversity in its animal diet.

In this research project, DNA metabarcoding was utilized to analyze fecal DNA from duck species wintering in a wetland complex in Hong Kong. By employing multiple markers, including *18S*, *trnL*, and *COI*, DNA metabarcoding offered a more comprehensive analysis of the diets of these species compared to DNA metabarcoding using fewer markers or traditional techniques like microscopy (da Silva et al., 2019). This multi-marker approach is particularly advantageous for animals with complex dietary habits, such as omnivores that consume plants and animals. Unlike other markers that target specific taxonomic groups, the *18S* marker plays a crucial role in providing insights into the relative proportions of food items spanning different kingdoms and phyla (Huang et al., 2021, Huang et al., 2022, Wei et al., 2024) in the diets of the ducks. A study in Japan using DNA metabarcoding with *trnL* and *COI* markers also reveals the diets of several duck species during the winter period (Ando et al., 2023), which uncovered a high diversity of consumed plant and invertebrate taxa. For example, *A. crecca* in Japan was found to consume at least 15 plant species from 12 families. The majority of plant materials were from the Araceae, Nymphaeaceae, and Poaceae families but not from Asteraceae (83% RRA in this study) (Ando et al., 2023).

Additionally, the study reveals that *A. crecca* consumed at least 15 families of arthropods, mollusks, or rotifers. However, the genetic markers used in the study could not reveal the relative proportions of plant versus animal taxa in the diet of *A. crecca*. Therefore, the use of a universal marker in this study therefore provides valuable insights into the relative proportion of higher taxonomic groups, e.g., plant and animal taxa, and demonstrates some duck species, e.g., *A. crecca* in this study, to be more specialized than others.

Comparing our results with those of studies conducted globally, we found that the dietary compositions of these duck species varied to different degrees across geographic locations. This variation underscores the flexibility of these duck species in their diets and their adaptable foraging strategies likely contribute to the sustainability of their populations. For example, *A. crecca* in Hong Kong showed the highest consumption of streptophytes, accounting for 94% RRA. Upon further examination, it was discovered that the streptophytes consumed were mainly asters from the Asteraceae family and grasses from the Poaceae family. In the Great Salt Lake, Utah, USA, a study examining the intestinal tracts above the gizzards of co-occurring *A. crecca* and *S. clypeata* wintering ducks reveals that both species consumed similar food taxa. During winter, both species consume over 70% of animal matter (including brine shrimp cysts) while increasing their intake of plant materials during fall and spring (Roberts and Conover, 2014), which dramatically contrasts with the findings in Hong Kong. However, similar to the *A. crecca* in Hong Kong, *A. crecca* wintering in the Camargue, southern France, consumed a high proportion of plant seeds (>80%) and significantly fewer invertebrates (<16%) during fall and winter. The plant species they found in their gullets included seven species from Cyperaceae, Poaceae, Potamogetonaceae, Amaranthaceae, and green algae (Brochet et al., 2012), but not from Asteraceae (83% RRA in this study). Another study on the diets of wintering *A. crecca* in Kern, California, USA, which examined the contents of oesophagi, found that plant seeds accounted for about 62% of their diet during fall and winter, with the remainder being animal matter. *Anas crecca* consumed more than nine plant species in this region, mainly from Poaceae and Lythraceae (Euliss Jr and Harris, 1987). The diverse dietary compositions of wintering *A. crecca* across various geographic locations highlight the adaptability and flexible foraging strategies of *A. crecca*.

For *S. clypeata*, streptophytes, including species from Asteraceae and grass Poaceae, were major components of *S. clypeata*’s diet in Hong Kong, accounting for 36% RRA, along with various taxa. Additionally, it exhibited a higher consumption rate of arthropods (36% RRA) compared to local *A. crecca* and *M. penelope*. A study analyzing the gizzards of wintering *S. clypeata* from 12 states in the USA revealed different proportions of plant (66%) and animal (35%) contents (McAtee, 1922) compared to the results of this study.

The USA study identifies a diverse range of items, including at least eight species of gastropods, eight species of Dytiscidae (water beetles), eight species of ostracods, and 53 species of plants, such as from families Poaceae, Potamogetonaceae, and Boraginaceae, among others. However, in another study conducted in Texas, USA, *S. clypeata* were predominantly herbivorous (plant >93.7%), with the contents in esophagus mainly comprising at least 22 plant species from e.g., the Polygonaceae and Poaceae families, as well as some gastropod species (Collins et al., 2017), in contrast to the Poaceae, Astraceae, and Acanthaceae families, and arthropods (>36% in *18S*) observed in our study. Furthermore, a study examining the contents of gizzards and gullets of *S. clypeata* wintering in Lake Tonga in Algeria reveals that their diets consisted entirely of plant materials without any inclusion of animal matter (Ayaichia et al., 2018). The study reported seven plant species, and the ducks predominantly consumed those from Typhaceae, Cyperaceae, Haloragaceae, and Ceratophyllaceae families (Ayaichia et al., 2018). In a similar study conducted in South Texas, a notable difference was observed in the dietary compositions of *S. clypeata* between freshwater and saltwater habitats. The study reveals that more animal matter was consumed in saltwater habitats (over 80%) than in freshwater habitats (50%). The researchers identified animal components in their esophagus and proventriculus that belonged to seven orders and representatives from one phylum, three classes, two families, and one genus. Notably, the primary animal matter consumed in saltwater environments included ostracods, foraminiferans, gastropods, and copepods (Tietje and Teer, 1996). The diets of *S. clypeata* examined in the study conducted in Japan, similarly using DNA metabarcoding, showed a lower diversity than our findings. The study indicated that the primary food sources for *S. clypeata* included various species of non-biting midges, mosquitoes, as well as plants from the Nelumbonaceae and Araceae families, which was distinct from those in Hong Kong (Ando et al., 2023).

Although research on the dietary compositions of *M. penelope* is limited, the dietary variations among *M. penelope* across different geographical locations have also been noted. Plant matter contributed a great proportion (40% RRA) to the diet of *M. penelope* wintering in Hong Kong; the plant matter is primarily composed of species from the Poaceae, Asteraceae, and families in Myrtales. Arthropods, including insects and malacostracans, accounted for 77% RRA within invertebrates consumed. In Vejlerne, Denmark, *M. penelope*’s diet mainly consisted of plant species from Poaceae, Juncaceae, Rosaceae, and Fabaceae, as analyzed using DNA metabarcoding (Svendsen et al., 2023). In Japan, wintering *M. penelope* mainly consumed plant species in the family Nelumbonaceae, Araceae, Ranunculaceae, Apiaceae, and Poaceae (Ando et al., 2023). The diverse dietary compositions of wintering *S. clypeata* and *M. penelope* in various geographic locations also emphasize the adaptability and flexible foraging tactics of these species. It is worth noting that comparing findings from dietary studies across different regions is challenging due to various factors. These factors include variations in the sampling seasons and digestive parts or materials examined, the taxonomic levels at which food items were identified, and differences in how studies analyzed their data and reported their results (Dessborn et al., 2011).

In addition to the interspecific dietary variability between different regions, our study further highlights the flexibility and adaptability of duck diets through the observed intraspecific dietary variations between individuals in the same habitat. While research on individual dietary variations among different duck species remains limited, our findings indicate that the overall and plant-based diets of *A. crecca* displayed much lower individual variability than those of *M. penelope* and *S. clypeata*, in terms of both taxa abundance and occurrence. The diet of most *A. crecca* individuals was dominated by a single taxon. In contrast, individuals of *M. penelope* and *S. clypeata* were highly variable in their dietary composition. These intraspecific dietary variations among duck individuals demonstrate their ability to adapt to diverse habitats and flexible food choices based on the availability of resources in the habitats. Such adaptability likely contributes to the widespread abundance of duck populations globally.

Previous studies on the foraging methods of the three duck species suggest that the foraging behaviors of *A. crecca* and *S. clypeata* are more similar to each other than to that of *M. penelope* (Klimas et al., 2022, Kooloos et al., 1989, Guillemain et al., 2002b). However, we observed a higher similarity between the diets of *S. clypeata* and *M. penelope*. The reason for this similarity therefore remains uncertain. *Anas crecca* generally forages via dabbling, upending, or grazing (Pöysä, 1987), primarily feeding at night during winter (Guillemain et al., 2002a). Previous studies have reported that the plant materials found in the digestive tracts of *A. crecca* consisted mainly of seeds (Olney, 1963), while vegetation shoots accounted for a limited proportion (Klimas et al., 2022). Limited research indicates that *A. crecca* exhibits selective feeding behavior, preferring small to medium-sized plant seeds (<4mm) and prey (Klimas et al., 2022). Previous studies in France revealed that *S. clypeata* and *M. penelope* foraged differently during winter, with *S. clypeata* engaging in dabbling or foraging deep in the water column by dipping and upending (Guillemain et al., 2000b, Guillemain et al., 2000a), while *M. penelope* predominantly grazed (Guillemain et al., 2002b). Similar to *A. crecca*, *S. clypeata* were mainly granivorous (Ayaichia et al., 2018). Previous research has proposed a sieving mechanism for *S. clypeata*, allowing it to filter and select food particles smaller than 4mm (Kooloos et al., 1989). According to previous studie, *S. clypeata* was observed foraging during both day and night time (Guillemain et al., 2000b, Guillemain et al., 2000a, Guillemain et al., 2002a). A study on the foraging behavior of *M. penelope*, it was found that *M. penelope* primarily grazes on green shoots (Mathers and Montgomery, 1997). During winter, *M. penelope* showed the highest peck rates on grass with a height of 30 mm, with peck rates decreasing on both taller and shorter grasslands (Durant and Fritz, 2006). Other studies observed that most *M. penelope* individuals engaged in water-dabbling for shoots during the observation period, with a small percentage involved in dig feeding, peck feeding, or upending depending on the tidal level (Mathers and Montgomery, 1996), and they mainly foraged in the daytime (von Känel, 1981).

While certain plant species like *Phragmites australis* identified in the diets of the duck species in this study have been documented as food sources for their conspecifics in other regions, our study also reveals the presence of other plant species not previously reported in studies of duck diets. For example, *Ficus* (figs) and *Kandelia obovata*, a kind of mangrove found in the Mai Po wetland, were consumed by the four duck species we studied. Furthermore, our research identified certain invertebrates that had not been previously documented in duck diets, such as *Metopograpsus frontalis* (crabs), *Paranais frici* (annelids), and *Tanytarsus formosanus* (non-biting midges).

Based on our research findings, we suggest that wet grasslands dominated by herbaceous plants, along with aquatic environments teeming with small aquatic invertebrates or zooplankton, serve as crucial foraging grounds for duck species wintering in Mai Po. To attract migratory duck species to winter in Mai Po, it is essential to focus on managing the Ramsar site and its surrounding areas. This includes maintaining or expanding pond areas and enhancing the abundance and diversity of herbaceous plant species in proximity to these ponds. Given that the studied duck species are primarily filter feeders and grazers, prioritizing grassland management, improvement, and restoration within the Ramsar site is key to promoting the growth of herbaceous plants, to ensure the availability of food resources for these duck species during their migration and maintain the health of their wintering habitats.

## Author contributions

Conceptualization - S.Y.W.S., Y.H.S., I.W.Y.S.; Methodology - S.Y.W.S., E.S.K.P.; Resources - S.Y.W.S., I.W.Y.S.; Investigation - E.S.K.P., L.Y.C., D.K.L., S.Y.W.S.; Formal analysis - P.Y.H.; Visualization - P.Y.H.; Writing - Original Draft - E.S.K.P., P.Y.H.; Writing - Review & Editing - E.S.K.P., P.Y.H., I.W.Y.S., Y.H.S., S.Y.W.S.; Supervision - S.Y.W.S.; Project administration - S.Y.W.S.; Funding acquisition - S.Y.W.S.

## Data Availability Statement

The datasets generated for this study can be found in the NCBI Sequence Read Archive (Accession numbers will be provided after the manuscript is accepted).

## Acknowledgments

We thank Hak-King Ying for his invaluable assistance with sample collection and Charis May Ngor Chan for her technical support. Special thanks to the World Wide Fund for Nature Hong Kong for permitting us to enter the Mai Po Nature Reserve. We would also like to acknowledge the Information Technology Services at the University of Hong Kong for providing the research computing facilities used for the computations in this study.

## Ethics approval statement

As this study does not involve live animals, permit is not required for this study.

## Permission to reproduce material from other sources

Illustrations of anatids used in figure 3 were reproduced with the permission of Lynx Edicions.

## Supplementary Materials and Methods

### DNA extraction and DNA barcoding to identify host species

DNA was extracted from fecal samples using E.Z.N.A. Tissue DNA Kit (Omega Bio-tek, Norcross, USA) following manufacturer’s protocol. We designed a pair of primers to specifically amplify a DNA minibarcode (142 bp) from the Folmer region of mitochondrial cytochrome c oxidase I (*COI*) gene, by aligning the *CO*I sequences from all locally occurring anatid species as well as other waterbird species that commonly winter in Hong Kong. PCRs were performed in 25 μl reactions comprising 5 μl of 5X GoTaq Flexi Buffer, 0.5 μl of 10 mM dNTP Mix, 3 μl of 25 mM MgCl2, 0.125 μl of 5 U/μl GoTaq G2 Flexi DNA Polymerase (all from Promega), 5 μl of extracted DNA, 1 μl of 10 μM forward primer (5’GCHATTAACTTCATYAC3’), 1 μl of 10 μM reverse primer (5’AAGAATGTGGTGTTTAGGTT3’), 2.5 μl of 10 % dimethyl sulfoxide (DMSO) (Sigma), 0.125 μl of 20 mg/ml bovine serum albumin (BSA) (New England Biolabs) and 6.75 μl of UltraPure DNase/RNase-Free Distilled Water (ultrapure water) (Invitrogen, Carlsbad, CA). PCRs were performed with the following condition: 95 °C for 2 min; 40 cycles of 95 °C for 30 sec, 45 °C for 30 sec, and 72 °C for 30 sec; and final extension at 72 °C for 5 min. All DNA minibarcodes obtained were sanger sequenced to confirm the identities of host species. Sequencing was carried out by the BGI (Shenzhen, China).

### Preparation of mock communities

We obtained 13 specimens of animal and plant species, including seven animal species belonging to the classes Malcostraca, Insecta, Polychaeta, and Actinopterygii as well as six plant species in the clades Rosids, Commelinids, and Monocots, from a local food market (Table S1). We extracted gDNA from the fresh tissues of animals and plants using the E.Z.N.A. Tissue DNA Kit (Omega Bio-tek, Norcross, USA) and the DNeasy Plant Pro Kit (Qiagen, Hilden, Germany), respectively. Six mock communities (MC1-MC6) were prepared according to Table S1, which shows the species compositions of each mock community. MC1-6 were prepared by mixing 12ng of gDNA from each of the included species.

### Preparation of DNA metabarcoding libraries through 2-step PCRs

For *18S* rDNA libraries, 1^st^ step PCRs were carried out in 30 μl reactions containing 6 μl of 5X GoTaq Flexi Buffer, 0.6 μl of 10 mM dNTP Mix, 3.6 μl of 25 mM MgCl_2_, 0.15 μl of 5 U/μl GoTaq G2 Flexi DNA Polymerase, 5 μl of extracted DNA, 0.6 μl of each assigned 10 μM forward and reverse primer uniquely tagged with heterogeneity spacer (Cruaud et al. 2017), 3 μl of 10 % DMSO (Sigma), 0.15 μl of 20 mg/ml BSA (NEB) and ultrapure water. Thermal cycling condition was 95 °C for 2 min; 25 cycles of 95 °C for 30 sec, 50 °C for 30 sec and 72 °C for 30 sec; and final extension at 72 °C for 5 min. For *COI* libraries, the 1^st^ step PCRs were carried out in 30 μl reactions containing 6 μl of 5X GoTaq Flexi Buffer, 0.6 μl of 10 mM dNTP Mix, 3.6 μl of 25 mM MgCl_2_, 0.15 μl of 5 U/μl GoTaq G2 Flexi DNA Polymerase, 4 μl of extracted DNA, 0.6 μl of each assigned 10 μM forward and reverse primer uniquely tagged with heterogeneity spacer (Cruaud et al. 2017), 3 μl of 10 % DMSO (Sigma), 0.15 μl of 20 mg/ml BSA (NEB), 6 μl of 10μM PNA blocker (Panagene), and ultrapure water. Blockers were used in *M. penelope*, *S. clypeata*, *A. crecca* samples except for *A. acuta* because the sample size of *A. acuta* (n=3) was very small. Thermal cycling conditions were 95°C for 2 min; 25 cycles of 95 °C for 30 s and 47 °C for 45 s, and final extension at 50 °C for 15 min. For *trn*L libraries, the 1^st^ step PCRs were carried out in 30 μl reactions containing 6 μl of 5X GoTaq

Flexi Buffer, 0.6 μl of 10 mM dNTP Mix, 3.6 μl of 25 mM MgCl_2_, 0.15 μl of 5 U/μl GoTaq G2 Flexi DNA Polymerase, 5 μl of extracted DNA, 0.6 μl of each assigned 10 μM forward and reverse primer uniquely tagged with heterogeneity spacer (Cruaud et al. 2017), 3 μl of 10 % DMSO (Sigma), 0.15 μl of 20 mg/ml BSA (NEB) and ultrapure water. Thermal cycling condition was 95 °C for 2 min; 25 cycles of 95 °C for 30 sec, 50 °C for 30 sec and 72 °C for 30 sec; and final extension at 72 °C for 5 min. We used the E.Z.N.A. Gel Extraction Kit (Omega Bio-tek) to clean up all PCR products generated from the two-step PCRs. Each of the 1^st^ step PCR products was eluted in 20 μl elution buffer and used as the templates for the 2^nd^ step PCRs. The 2^nd^ step PCRs of both markers were performed with the same conditions: 2^nd^ step PCRs were performed in 45 μl reactions comprising 9 μl of 5X GoTaq Flexi Buffer, 0.9 μl of 10 mM dNTP Mix, 5.4 μl of 25 mM MgCl_2_, 0.225 μl of 5 U/μl GoTaq G2 Flexi DNA Polymerase, 20 μl (*18S* and *trnL*) and 18.5 μl (*COI*) of 1^st^ step PCR products, 0.9 μl of each assigned 10 μM forward and reverse index primers, 4.5 μl of 10 % DMSO (Sigma) and ultrapure water. PCR condition for *18S* and *trnL* was 95 °C for 2 min; 95 °C for 30 sec, 55 °C for 30 sec, and 72 °C for 30 sec (15 cycles for *18S* and 20 cycles for *trnL*); and final extension at 72 °C for 5 min. PCR condition for *COI* was 95 °C for 2 min; 15 cycles of 95 °C for 30 sec, 55 °C for 45 sec; and final extension at 55 °C for 15 min.

## Supplementary Figures

**Figure S1.**
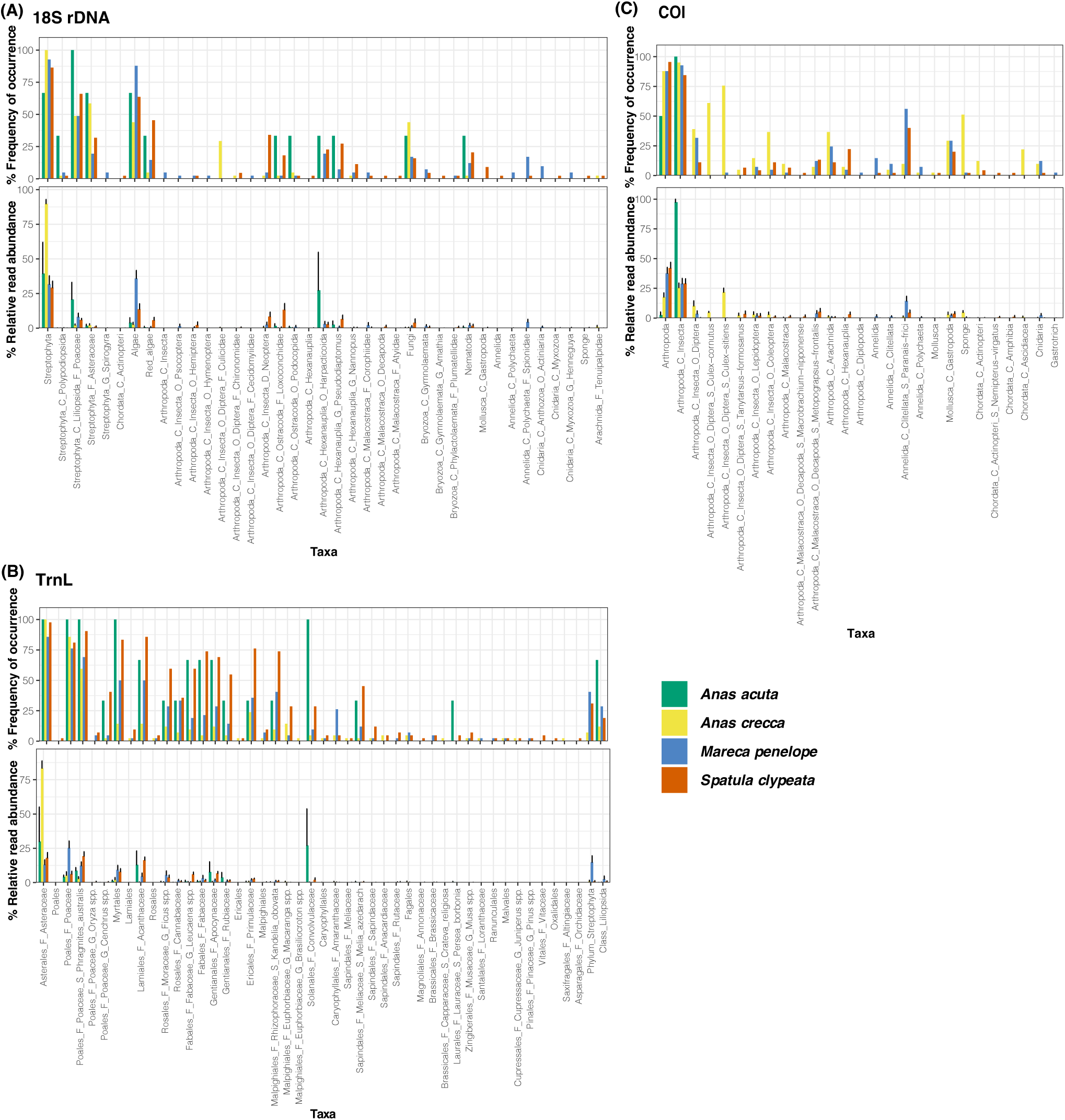
Occurrences and abundances of dietary taxa detected in the fecal samples of anatids using (A) *18S* rDNA (n=129), (B) *trnL* (n=129), and (C) *COI* markers (n=129). Frequency of occurrence, means and standard errors of the relative read abundance of each taxon, were presented. Class, C; Order, O; Family, F; and Genus, G.

**Figure S2.**
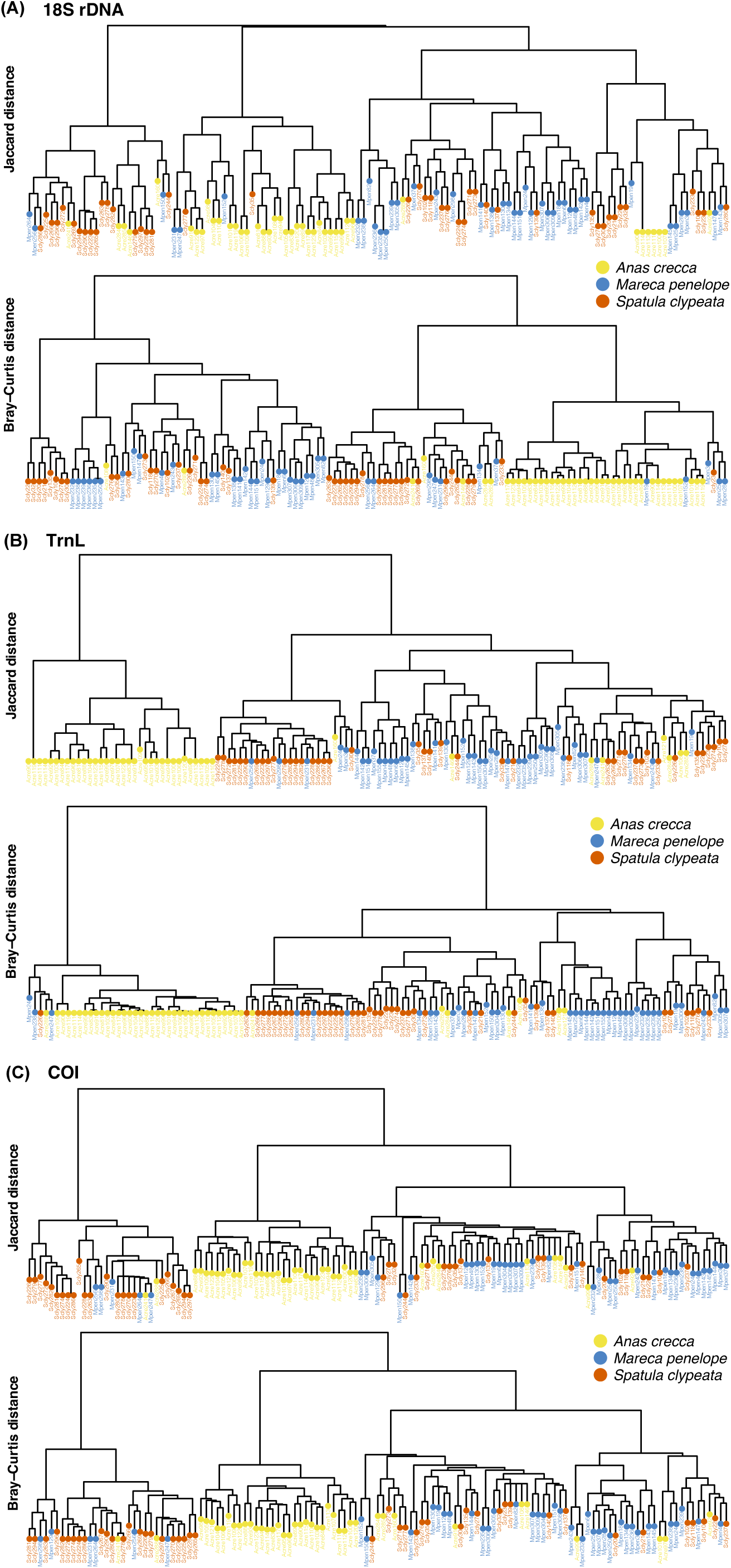
Hierarchical clustering dendrograms showed the differences in dietary compositions observed among anatid individuals, as identified through the use of (A) *18S*, (B) *trnL*, and (C) *COI* markers. We analyzed fecal samples from *Spatula clypeata* (n=44 for *18S*, n=42 for *trnL*, and n=45 for *COI*), *Mareca penelope* (n=41 for *18S* and *COI*; n=42 for *trnL*), and *Anas crecca* (n=41 for *18S* and *COI*; n=42 for *trnL*).

